# PiggyBac Transposase Mediated Inducible Trophoblast-specific Knockdown of Mechanistic Target of Rapamycin in Mice Decreases Placental Nutrient Transport and Inhibits Fetal Growth

**DOI:** 10.1101/2024.02.20.581180

**Authors:** Fredrick J Rosario, Johann Urschitz, Haide Razavy, Marlee Elston, Powell L Theresa, Thomas Jansson

## Abstract

Abnormal fetal growth is associated with perinatal complications and adult disease. The placental mechanistic target of rapamycin (mTOR) signaling activity is positively correlated to placental nutrient transport and fetal growth. However, if this association represents a mechanistic link, it remains unknown. We hypothesized that trophoblast-specific mTOR knockdown in late pregnant mice decreases placental nutrient transport and inhibits fetal growth. PiggyBac Transposase-Enhanced Pronuclear Injection was performed to generate transgenic mice containing a trophoblast-specific Cyp19I.1 promoter-driven, doxycycline-inducible luciferase reporter transgene with a *Mtor* shRNAmir sequence in its 3’ untranslated region (UTR). We induced mTOR knockdown by administration of doxycycline starting at E14.5. Dams were euthanized at E 17.5, and trophoblast-specific gene targeting was confirmed. Placental mTOR protein expression was reduced by ∼68% in these animals, which was associated with a marked inhibition of mTORC1 and mTORC2 signaling activity. Moreover, we observed a decreased expression of System A amino acid transporter isoform SNAT2 and the System L amino acid transporter isoform LAT1 in isolated trophoblast plasma membranes and lower fetal but not placental weight. Inhibition of trophoblast mTOR signaling in late pregnancy is mechanistically linked to decreased placental nutrient transport and reduced fetal growth. Modulating placental mTOR signaling may represent a novel intervention in pregnancies with abnormal fetal growth.

## Introduction

Abnormal fetal growth, i.e., intrauterine growth restriction (IUGR) or fetal overgrowth, is associated with increased perinatal morbidity and mortality and is strongly linked to the development of metabolic and cardiovascular disease in childhood and later in life ^1–6^. Fetal growth is largely determined by oxygen and nutrient availability, controlled by placental blood flow and nutrient transport capacity^7,8^. Emerging evidence suggests that changes in placental amino acid transport expression and activity may contribute to abnormal fetal growth. Specifically, the activity of System A and L amino acid transporters has been reported to be decreased in human IUGR ^9–12^. In addition, various studies have reported an upregulation of placental System A and L transport activity in fetal overgrowth^13–15^. In primate and rodent models of IUGR, down-regulation of placental System A and L transporter activity precedes fetal growth restriction^16–18^, consistent with the possibility that altered placental amino acid transport is mechanistically linked to abnormal fetal growth. Furthermore, changes in fetal nutrient availability program the fetus for metabolic and cardiovascular disease later in life^3,19–21^.

The System L transporter is a sodium-independent exchanger mediating the transport of large neutral, predominantly essential amino acids, including leucine. System L is a heterodimer composed of a light chain, commonly known as LAT1 (*SLC7A5*) or LAT2 (*SLC7A8*), and a heavy chain, referred to as 4F2hc/CD98 (*SLC3A2*)^22^. Sodium-dependent neutral amino acid transporter 1 (SNAT1) (*SLC38A1*), SNAT2 (*SLC38A2*), and SNAT4 (*SLC38A4*) isoforms of the System A transporter are all expressed in the placenta^23^ and preferentially mediate the active uptake of a range of neutral non-essential amino acids, including alanine, serine, and cysteine^24^.

The mechanistic target of the rapamycin (mTOR) signaling pathway serves as a key regulator of many different cellular functions, ranging from cellular metabolism to growth and survival^25^. mTOR responds to various signals, such as hormones, growth factors, nutrients, energy availability, and stress signals^26,27^. mTOR is present in two distinct complexes: mTOR complex 1 (mTORC1) and mTORC2^28,29^. The trophoblast mTOR pathway is considered a master regulator of placental function. For example, studies in cultured primary human trophoblast cells have demonstrated that mTOR signaling is a positive regulator of trophoblast amino acid (System A and L), folate transport, and mitochondrial respiration ^30–32^. In addition, mTOR regulates glucose uptake in trophoblast-derived cell line cultures ^33^. Both mTORC1 and mTORC2 signaling regulate trophoblast expression of various genes involved in ribosome biosynthesis, inflammation, micronutrient transport, and angiogenesis, representing novel links between mTOR signaling and multiple placental functions necessary for fetal growth and development ^34,35^. However, data unequivocally demonstrating that changes in trophoblast mTOR signaling regulate placental function *in vivo* are lacking.

Placental mTOR signaling has been proposed to serve as a critical hub for homeostatic control of fetal growth in response to maternal nutrition and metabolism perturbations, as well as uteroplacental blood flow ^36,37^. In line with this model, we and others demonstrated that placental mTOR signaling is inhibited in IUGR in women ^12,38–41^and a range of animal models^17,18,42–48^, while it is activated in fetal overgrowth^14,49–51^. Furthermore, placental mTOR signaling is reduced before the reductions in fetal growth, indicating that changes in placental mTOR signaling could be the cause rather than a consequence of abnormal fetal growth^18^. Interestingly, inhibition of placental mTOR decreases the weight of female but not male fetuses^52^. However, compelling mechanistic evidence for the key role of trophoblast mTOR in regulating placental function and fetal growth is lacking.

Homozygous mTOR knockout embryos die shortly after implantation due to impaired cell proliferation in both embryonic and extraembryonic compartments ^53^, and maternal administration of the mTORC1 inhibitor rapamycin to pregnant mice results in IUGR ^54^. Hence, these results preclude defining the specific role of trophoblast mTOR signaling in regulating placental function and fetal growth. We hypothesized that trophoblast-specific mTOR knockdown in late pregnant mice decreases placental nutrient transport and inhibits fetal growth. We employed *piggyBac* transposase-enhanced transgenesis^55^ to achieve trophoblast-specific gene modulation. Specifically, PiggyBac Transposase-Enhanced Pronuclear Injection was performed to generate transgenic mice by injecting vectors containing a trophoblast-specific Cyp19I.1 promoter-driven, doxycycline-inducible luciferase reporter transgene with mTOR shRNAmir sequence in its 3’ untranslated region (UTR), into one-cell B6D2F2 embryos. Here we report that a reduction in trophoblast mTOR signaling is mechanistically linked to decreased placental nutrient transport and reduced fetal growth. We propose that placenta-specific targeting of mTOR signaling represents a promising avenue to improve outcomes in pregnancies complicated by abnormal fetal growth.

## Materials and Methods

### Animals and Diet

The Institutional Animal Care and Use Committee (Protocol # 13-1697) of the University of Hawaii approved all protocols conducted per NIH guidelines. All mice had *ad libitum* access to food and water.

### Construction of plasmid DNA

We used the Broad Institute’s Gene Perturbation website to identify five different shRNAs with *Mtor* specific target sequences. Plasmids containing these shRNAs were obtained from Sigma (St. Louis, MO). We determined that the targeting sequence CCGTCCCTACATGGATGAAAT was most efficient in knocking down mTOR in primary human trophoblast cells. The basis for the final plasmid used for transgenesis was a commercially obtained TET-ON micro-RNA (miR30) adapted shRNA vector targeting the RNA of the mouse glucose transporter 1 (*Glut1,* V3THS_321626, Open Biosystems). This plasmid-encoded turboRFP under a TRE-3G promoter, with the reverse tetracycline-controlled transactivator 3 (rTTA3) driven by a ubiquitin C gene (UBC) promoter. The shRNAmir was located in the 3’UTR of turboRFP. We replaced turboRFP with the luciferase reporter gene to enable *in vivo* assessment of transgene expression. The UBC promoter was replaced with the trophoblast-specific Cyp19I.1 promoter ^56,57^ to limit the luciferase and shRNAmir transgene expression to trophoblast cells. Finally, we exchanged the shRNA target sequence of *Glut1* with that of *Mtor*. These steps were all performed by restriction and insertion cloning. The transgene was then cloned into a pENTR1A vector (Thermo Fisher, Waltham, MA USA) to facilitate the final step, the recombination of this *Mtor* knockdown pENTR1A vector with our pmhyGENIE-3 piggyBac vector^55,58^, to generate the final construct (**Figure 1**). The correct constructs were expanded, purified with gravity-flow columns, and used for transgenesis experiments.

**Figure 1:**
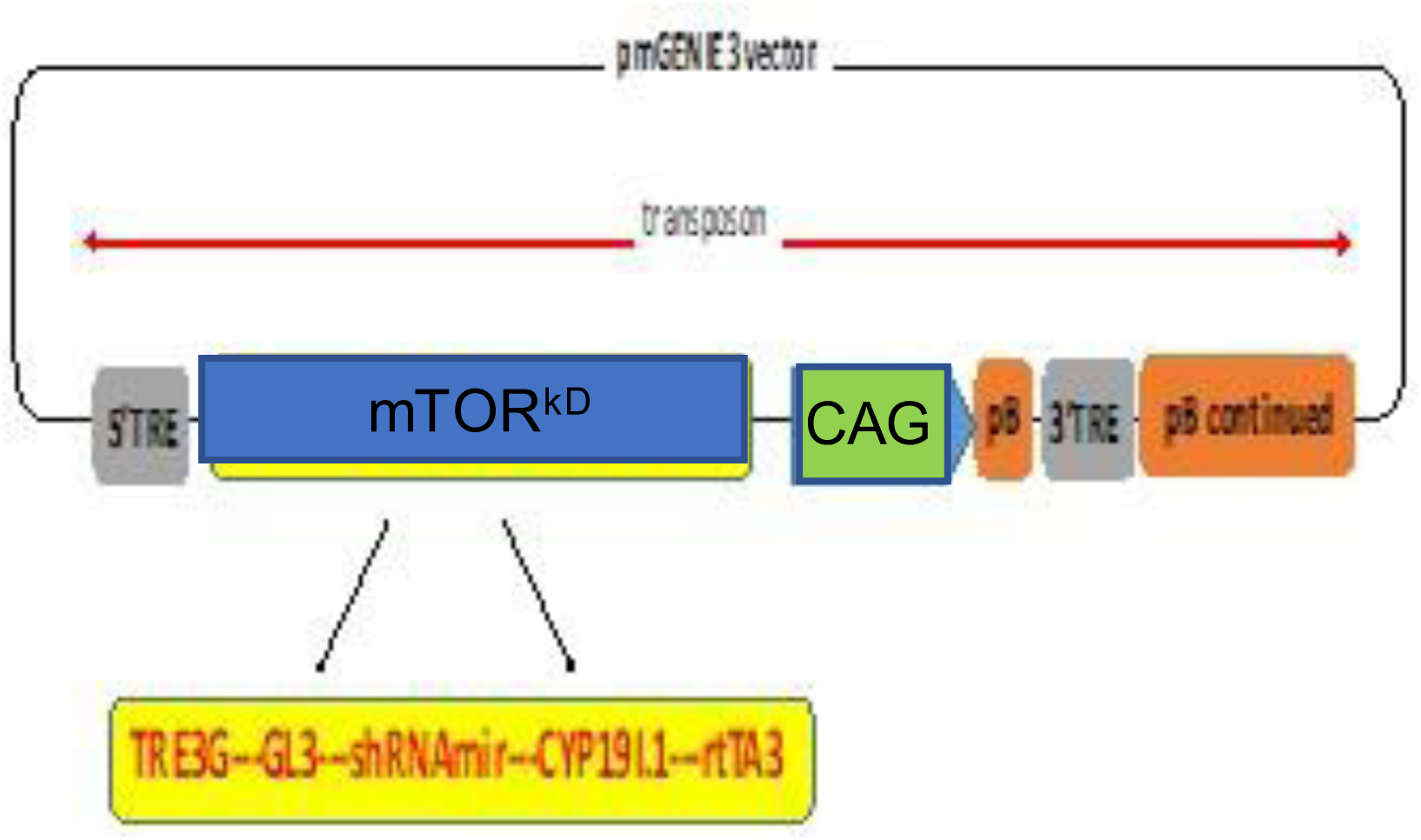
Transgene construct. A transgene construct is depicted, consisting of TRE3G & rtTA3, Tet-On system; GL3, luciferase reporter gene; shRNAmir, *Mtor* knockdown shRNAmir; CYP19I.1, Aromatase P450 fragment as the promoter for trophoblast-specific gene expression; CAG, CMV early enhancer/chicken beta-actin promoter; pB, piggyBac transposase and TRE, terminal repeat elements.

### Development of a trophoblast-specific inducible mTOR knockdown mouse

Transposase-enhanced pronuclear injection (te-PNI) was performed ^55,58–60^ using B6D2F1 once-cell-stage embryos (B57BL/6 x DBA/2) and CD1 mice purchased from Jackson Laboratories. Genomic DNA was isolated from pups, and genotyping PCR was performed (**Figure 2**). To determine the number of transgenes for the founder generation as well as F1 (crossed with WT B57BL/6), we performed transgene copy number assays by duplex Taqman real-time PCR, and results were analyzed using Applied Biosystems‘s Copy Caller software.

**Figure 2:**
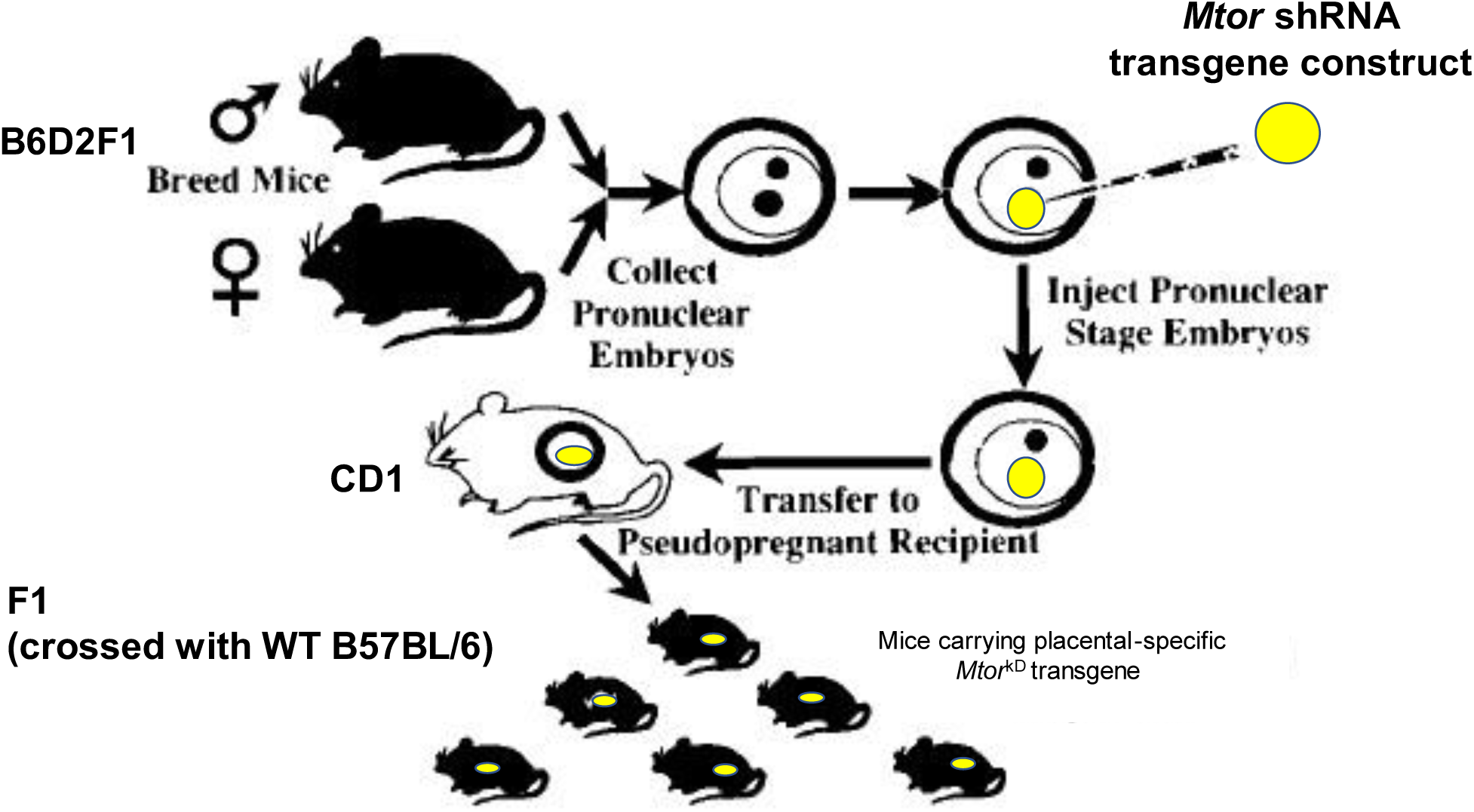
A schematic of the experimental design for developing a trophoblast-specific inducible *Mtor* knockdown mouse. Transposase-enhanced pronuclear injection was performed using B6D2F1 (B57BL/6 x DBA/2) and CD1 mice. Genomic DNA was isolated from pups, and genotyping PCR was performed to determine the number of transgenes for the founder generation and F1 (crossed with WT B57BL/6).

Trophoblast-specific *Mtor* knockdown (*Mtor*^kD^) was induced at E14.5 by administration of doxycycline 2.5 mg/kg (IP). Doxycycline demonstrated no potential to cause toxicity (data not shown). Transgenic animals in which vehicle was administered at E14.5 and wild-type females served as controls. At E17.5, there was no visible signal in transgenic animals in which *Mtor* knockdown had not been induced by doxycycline (**Figure 3**). In contrast, the luciferase signal could be detected in the placentas of dams that received doxycycline (**Figure 3**), with no visible signal in the embryo proper or maternal tissues.

**Figure 3:**
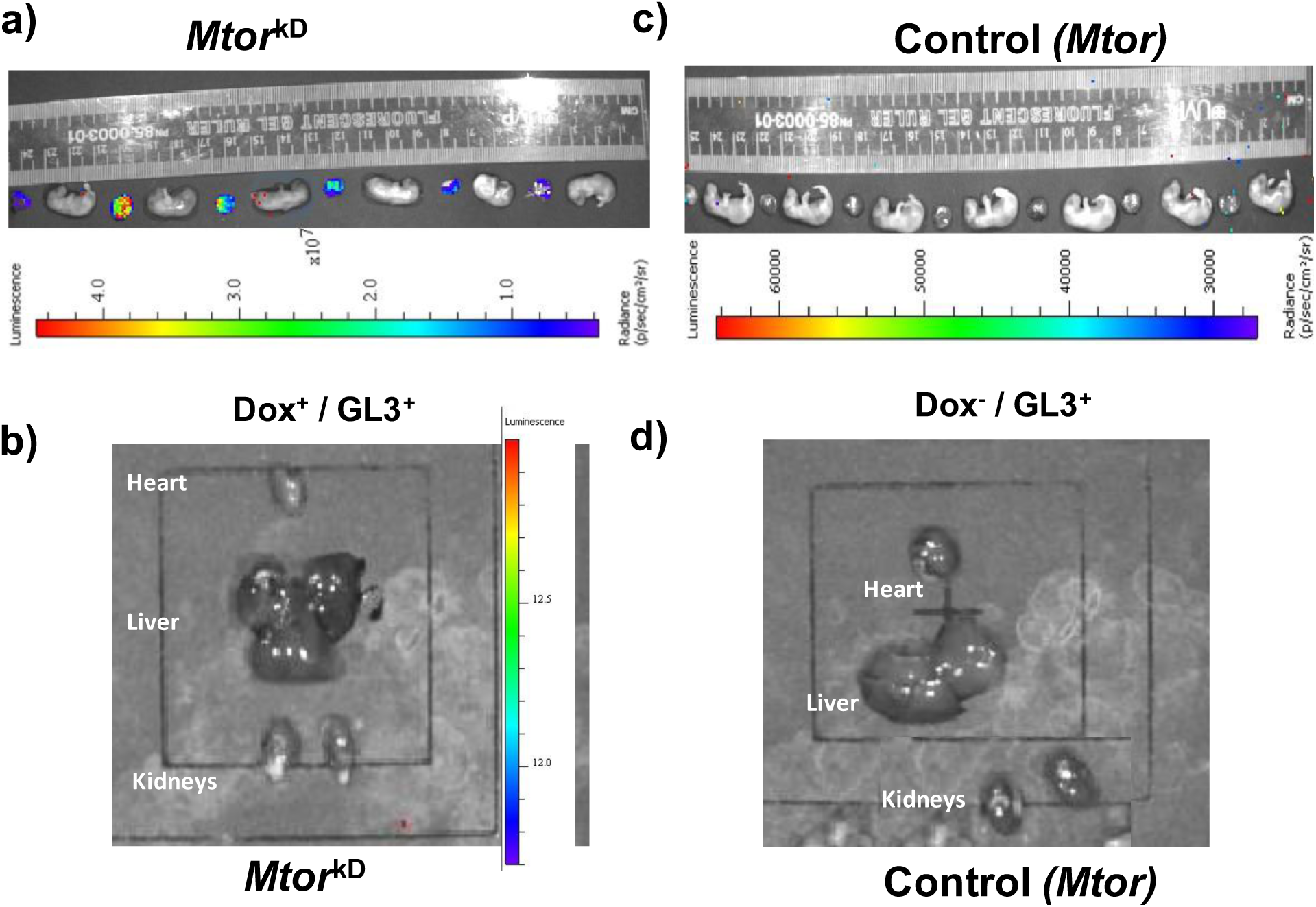
Placenta-specific inducible *Mtor* knockdown. *Mtor* knockdown was induced by the administration of doxycycline at E14.5 in pregnant mice transgenic for a construct including a Tet-On system, luciferase, *Mtor* shRNA mir, and CYP19I.1. The vehicle was administered in the control transgenic animals (No Doxycycline). Laparotomy was performed at E17.5, and the luciferase signal was detected in isolated placentas and fetuses using an in vivo imaging system (IVIS). The luciferase signal could be detected only in the (a) placentas of dams that received doxycycline with no visible signal in the embryo proper, (b) maternal tissues, or (c-d) in transgenic animals in which *Mtor* knockdown had not been induced by doxycycline.

### Collection of placental tissue

Both control and experimental group dams were euthanized at E17.5 by carbon dioxide inhalation. Following laparotomy, fetuses and placentas were collected and rapidly dried on blotting paper. After that, any remaining adherent fetal membranes were removed from fetuses and placentas before weighing. All placentas in each litter were pooled and washed briefly in PBS and transferred to 3 ml of ice-cold buffer D [250 mM sucrose, 10 mM HEPES-Tris, and 1 mM EDTA (pH 7.4) at 4°C]. A dilution of 1:1,000 protease and phosphatase inhibitor cocktail (Sigma-Aldrich, St. Louis, MO) was added to the solution, which was then homogenized using a Polytron (Kinematica, Bohemia, NY). The mixture was frozen in liquid nitrogen and stored at −80°C until further processing and analysis.

### Isolation of Trophoblast plasma membranes

The trophoblast plasma membrane (TPM) is the maternal-facing plasma membrane of syncytiotrophoblast layer II of the mouse placentas, which is believed to be functionally similar to the syncytiotrophoblast microvillous plasma membrane of the human placenta ^61^. As described elsewhere, TPMs were isolated from frozen placental homogenates using differential centrifugation and Mg2+ precipitation^61^. A Lowry assay (Bio-Rad, CA) was carried out to determine the protein concentration of TPM. The enrichment of TPM was assessed using the TPM/homogenate ratio of alkaline phosphatase activity. There was no significant difference in the average enrichment of alkaline phosphatase in TPM vesicles isolated from *Mtor* knockdown animals (17.1 ± 3.2, n=3) as compared to placentas of transgenic animals injected with vehicle (17.5 ± 6.5, n=3) or placentas of wild type animals (17.7± 1.9, n=3).

### Western blot analysis

Western blot analysis was carried out as described ^14,18,32,43^. Briefly, 10-15 μg of placental homogenates or 5-10 μg of TPM were loaded onto a NuPAGE Bis-Tris Gels (Thermo Fisher Scientific), and electrophoresis was performed using MOPS-SDS running buffer at a constant 100 V for 1 h. After gel electrophoresis, the separated proteins were transferred onto nitrocellulose membranes using NuPAGE™ Transfer Buffer (Invitrogen) at a constant 30 V for 1 hr. After transfer, nitrocellulose membranes were blocked in 5% blotting grade blocker nonfat dry milk (Bio-Rad, Hercules, CA) in TBS (wt/vol) plus 0.1% Tween 20 (vol/vol) for overnight at 4°C. Following serial washing in TBS plus 0.1% Tween 20 for 1 h at room temperature, the membrane was incubated with appropriate primary antibody overnight at 4°C. Subsequently, the membranes were washed in TBS plus 0.1% Tween 20 for 1 h at room temperature, followed by incubation for one hour with the corresponding secondary peroxidase-labeled antibodies. Following the washing step in TBS plus 0.1% Tween 20, the bands were visualized using enhanced chemiluminescence detection reagents from Thermo Fisher Scientific (Waltham, MA, USA). Blots were stripped ^14,18,43^ and reprobed for β-actin. Image J software from the National Institutes of Health (Bethesda, MD) was used to perform densitometry analysis of the Immunoblots. The mean density of the control sample bands was given a value of 1 for each protein target, and the data is presented in relation to the control. The relative density of the target protein in each lane was normalized by dividing it by the density of the corresponding β-actin band, which served as a loading control. Additionally, no significant difference was observed in β-actin expression between the control and experimental groups (data not presented).

Isolated TPM was used to measure the protein expression of System A amino acid transporter isoform (SNAT) 2 *(SLC38A2)* and the System L amino acid transporter isoform LAT1(*SLC7A5*). A polyclonal SNAT2 antibody generated in rabbits was received as a generous gift from Dr. V. Ganapathy and Dr. P. Prasad at the University of Georgia, Augusta. Antibodies targeting the LAT1 were produced in rabbits and received as a generous gift from Dr Yoshikatsu Kanai from Osaka University, Osaka, Japan. The specificity of SNAT2 and LAT1 antibodies has previously been validated using gene-silencing techniques ^62,63^. Furthermore, mTOR signaling pathway downstream signaling activity in placental homogenates was also measured by Western blot for total and phosphorylated forms of mTOR, S6 (Ser-235/236), 4EBP1 (Thr-70) and SGK1 (Ser-422), Akt (Ser-473).

### TPM amino acid transporter activity measurements

The TPM System A and L amino acid transporter activity was measured using radiolabeled amino acids and rapid filtration techniques^43,61^. TPM vehicles from WT, *Mtor*, and *Mtor*^kD^ groups were preloaded by incubation in 300 mM mannitol and 10 mM Hepes-Tris, pH 7.4 overnight at 4°C. Subsequently, TPM vesicles were resuspended in a small volume of the same buffer after being pelleted (final protein concentration: 5–10 mg ml^-1^). Membrane vesicles were kept on ice until transport activity measurements were performed. Immediately before transport activity measurements, TPM vesicles were warmed to 37°C. At time zero, 30 μl TPM vesicles were quickly mixed (1:2) with the incubation buffer containing [^14^C] methyl-aminoisobutyric acid (MeAIB, 150 μM) with or without Na^+^ or L-[^3^H] leucine (0.375 μM). Based on previous time course studies^43^, uptake at 15 seconds was used in all subsequent experiments. The uptake of radiolabeled substrate was terminated by adding 2 ml of ice-cold PBS. Subsequently, vesicles were rapidly separated from the substrate medium by filtration on mixed ester filters (0.45 μm pore size, Millipore Corporation, Bedford, MA, USA) and washed with 3 × 2 ml of PBS. Each condition was studied in duplicate for each membrane vesicle preparation in all uptake experiments. Filters were dissolved in 2 ml liquid scintillation fluid (Filter Count, PerkinElmer, Waltham, MA, USA) and counted. Appropriate blanks were subtracted from counts and uptakes expressed as pmol (mg protein) ^−1^. Na^+^-dependent uptake of MeAIB (corresponding to system A activity) was calculated by subtracting Na^+^-independent from total uptakes. Mediated uptake was calculated for leucine by subtracting non-mediated transport, as determined in the presence of 20 mM unlabeled leucine, from total uptake.

### Data presentation and statistics

Data are presented as mean ± SEM. Statistical significance between WT, *Mtor*, and *Mtor*^kD^ was determined by one-way ANOVA with Tukey–Kramer multiple comparisons post hoc test (P < 0.05).

## Results

### Trophoblast-specific inducible *Mtor* knockout decreases fetal weight in mice

As shown in **Figure 4a**, trophoblast-specific inducible *Mtor* knockout decreases fetal weight (p=0.02, n=3-4 dams/each group) as compared to WT, and *Mtor* (Control) groups. However, placental weights were comparable between groups (**Figure 4b**).

**Figure 4:**
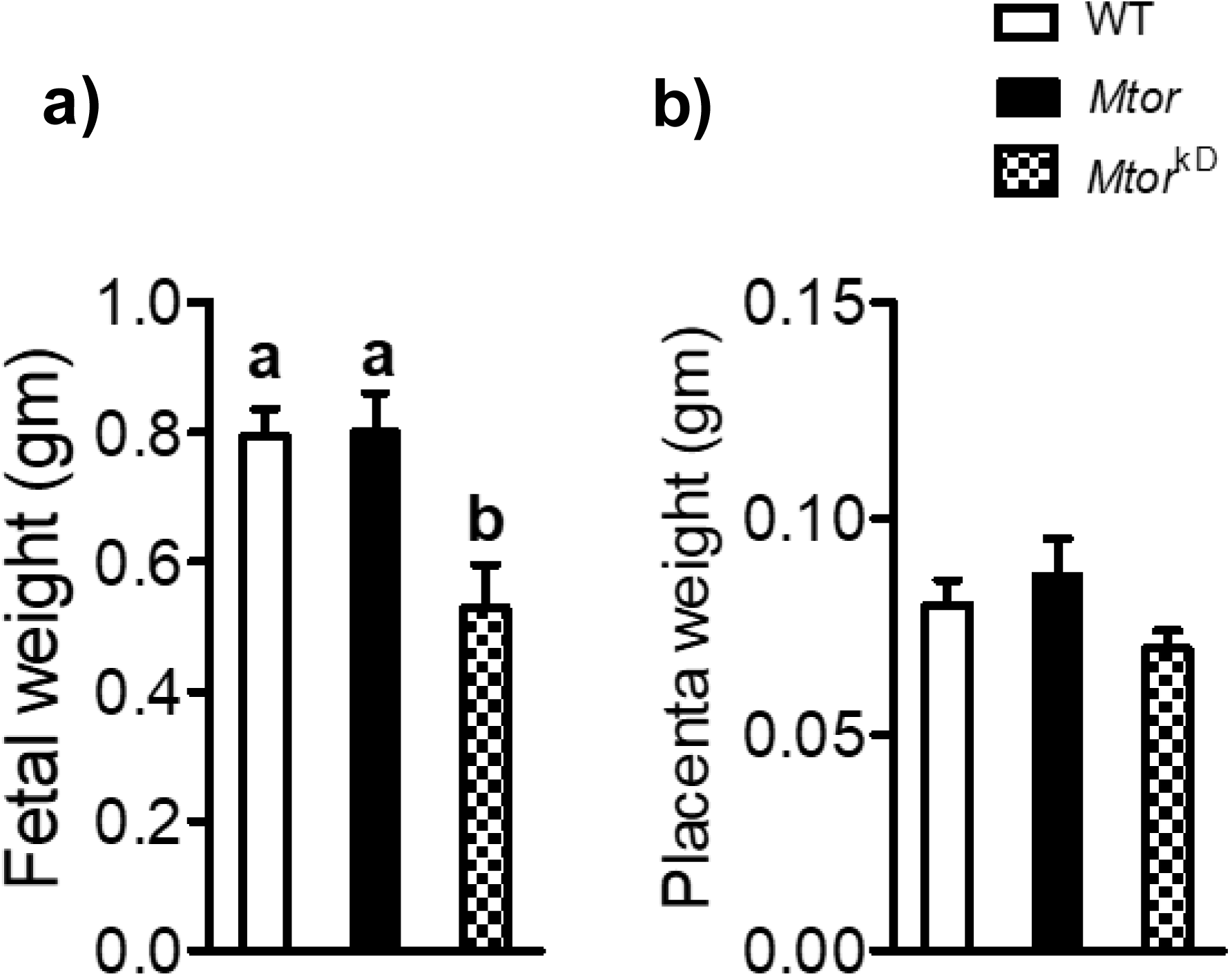
Fetal and placental weights. Compared to control and wild-type mice, trophoblast-specific *Mtor* knockdown (induced by doxycycline at E14.5) mice exhibit reduced fetal weights but not placental weights at embryonic day 17.5. Values are expressed as means ± SEM. Means without a common letter are statistically different by one-way ANOVA with Tukey–Kramer multiple comparisons post hoc test (P < 0.05).

### Trophoblast-specific inducible *Mtor* knockdown decreases placental mTORC1 signaling

To further explore the mechanisms linking trophoblast-specific inducible *Mtor* knockdown to decreased fetal growth, we used Western blot to determine functional readouts of the *Mtor* signaling pathways in WT, *Mtor*, and *Mtor*^kD^ placentas. Placental mTOR protein expression was 68 % (p=0.009, n=3-4 dams/each group) lower in the *Mtor*^kD^ group than in the WT and *Mtor* (Control) groups (**Figure 5 a-b**). Placental-specific *Mtor* knockdown significantly decreased 46 % (p=0.01, n=3-4 dams/each group) the S6 ribosomal protein phosphorylation at Serine 235/236, a *mTORC1* downstream target. Total S6 ribosomal protein expression level was comparable between groups (**Figure 5b**). Furthermore, trophoblast-specific knockdown of *Mtor* decreased 66 % (p=0.0001, n=3-4 dams/each group) the phosphorylation of 4E-BP1 (Threonine-70) in the *Mtor*^kD^ group as compared to the WT and *Mtor (*Control) groups (**Figure 5 a-b**). However, placental-specific knockdown of *Mtor* increased the total 4E-BP1 expression (+95%, p=0.006, n=3-4 dams/each group) compared to the WT and *Mtor* (Control) groups at E 17.5.

**Figure 5:**
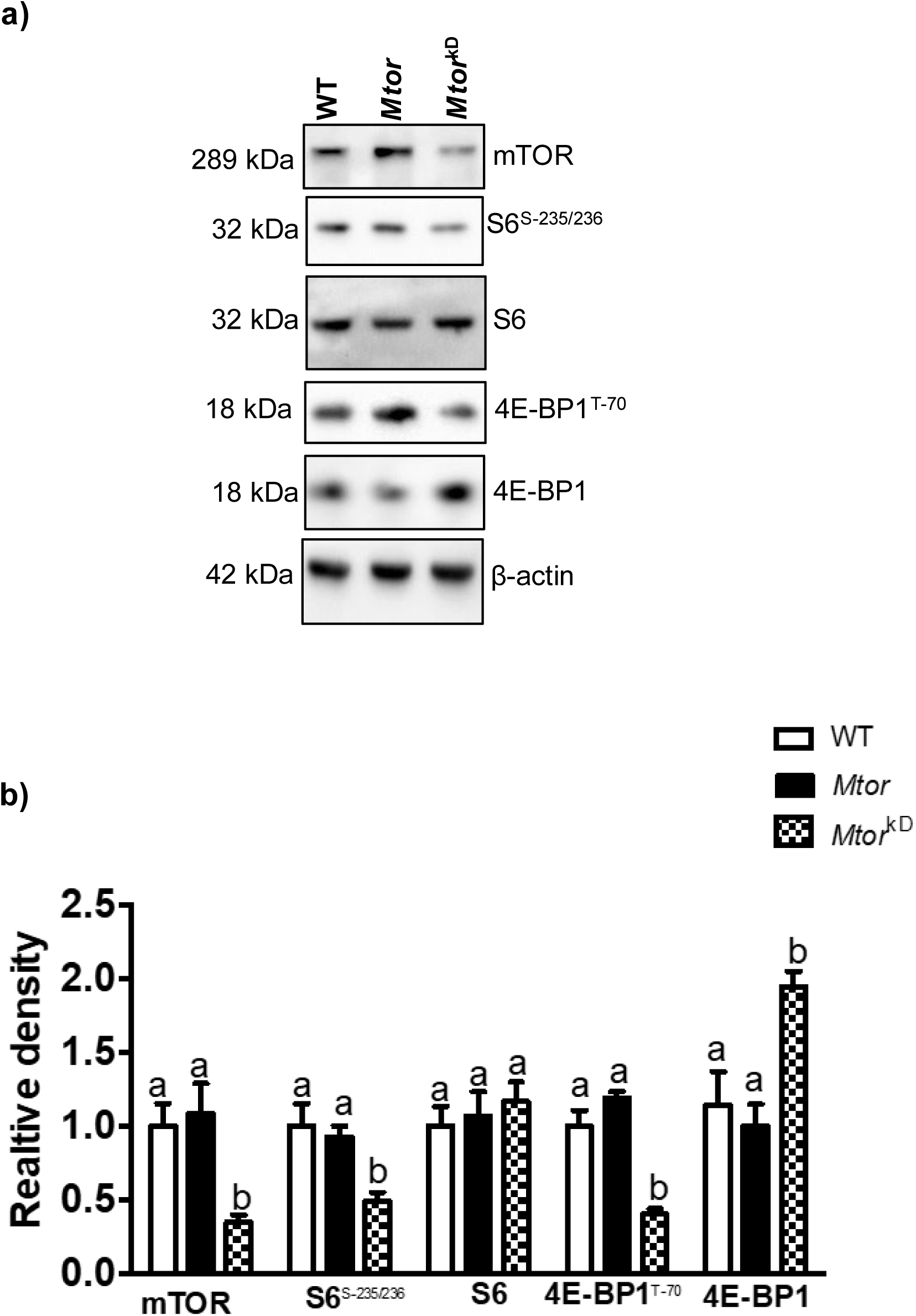
Inhibition of placental mTORC1 signaling following induction of trophoblast-specific *Mtor* knockdown. Trophoblast-specific mTOR knockdown was induced by the administration of doxycycline starting at E14.5 (*Mtor*^kD^). The vehicle was administered at E14.5 in control animals, resulting in mice without mTOR knockdown (*Mtor*). At E17.5, animals were sacrificed, and placentas from each litter were pooled and homogenized. Western blot determined the total expression and phosphorylation of key intermediates in the mTORC1 signaling pathways. (a) Representative western blots of mTOR, S6^Serine-235/236^, total S6, 4E-BP1 ^Threonine-70^, and 4E-BP1 expression in placental homogenates of WT, *Mtor* and *Mtor*^kD^ placentas. Equal loading was performed. (b) Summary of the western blot data. n = 3-4 dams in each group. WT = wild type. Values are expressed as means ± SEM. Means without a common letter are statistically different by one-way ANOVA with Tukey–Kramer multiple comparisons post hoc test (P < 0.05).

### Trophoblast-specific inducible *Mtor* knockdown decreases placental mTORC2 signaling

Placental-specific *Mtor* knockdown significantly decreased the phosphorylation of SGK1 at Serine-422 (−48%, p=0.001, n=3-4 dams/each group) and protein kinase B (Akt-Ser 473; −52%, p=0.004, n=3-4 dams/each group), functional readouts of mTORC2 activity, as compared to WT and *Mtor* (Control) group at E 17.5 (**Figure 6 a-b**). Moreover, the total expression of SGK1 and Akt was not different in WT, *Mtor*, and *Mtor*^kD^ groups.

**Figure 6:**
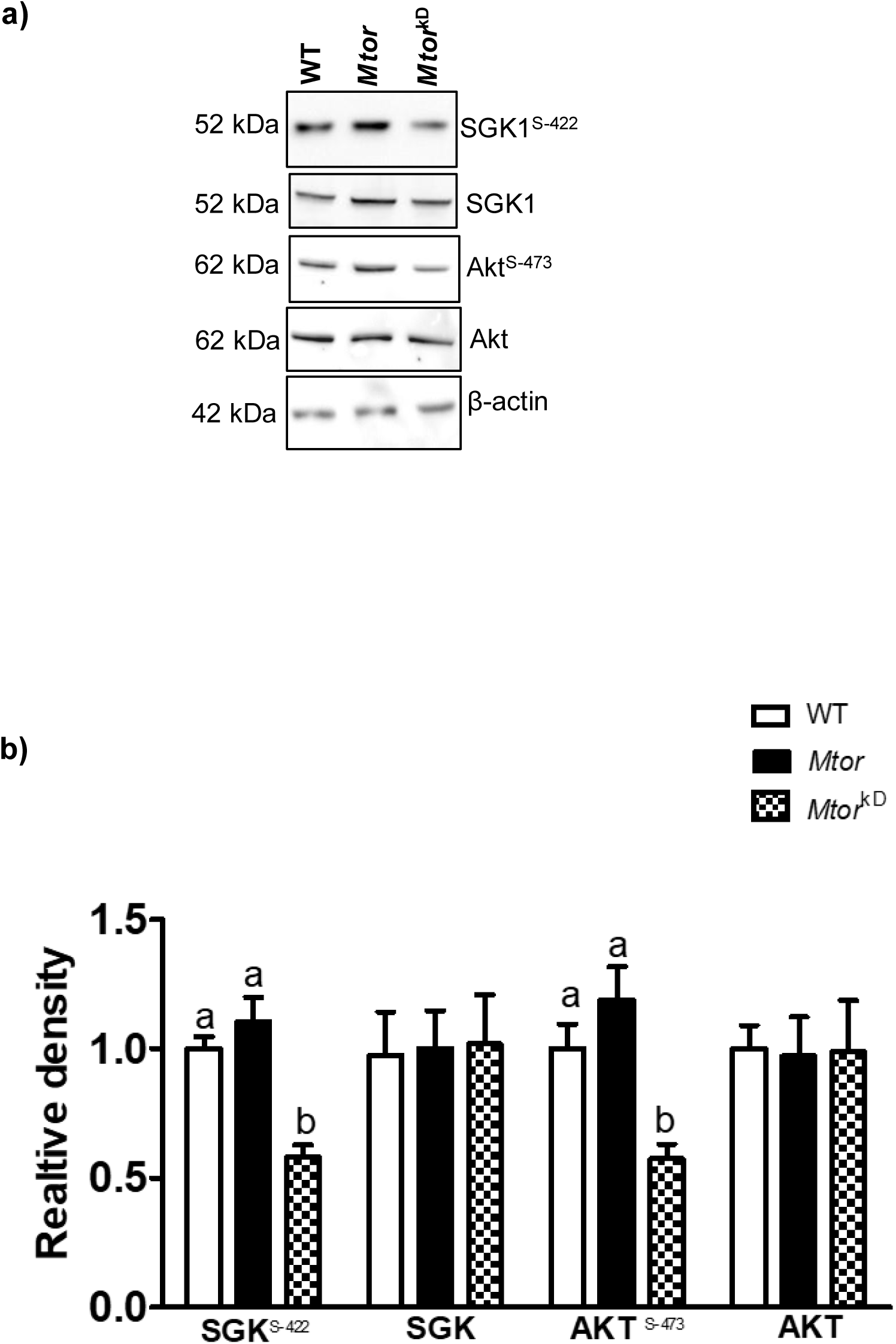
Inhibition of placental mTORC2 signaling following induction of trophoblast-specific *Mtor* knockdown. Trophoblast-specific mTOR knockdown was induced by the administration of doxycycline starting at E14.5 (*Mtor*^kD^). In control animals, the vehicle was administered at E14.5, resulting in mice without mTOR knockdown (mTOR). At E17.5, animals were sacrificed, and placentas from each litter were pooled and homogenized. Western blot determined the total expression and phosphorylation of key intermediates in the mTORC2 signaling pathways. (a) Representative western blots of SGK1^Serine-422^, total SGK1, Akt ^Serine-473^, and Akt expression in placental homogenates of WT, *Mtor,* and *Mtor*^kD^ placentas. Equal loading was performed. (b) Summary of the western blot data. n = 3-4 dams in each group. WT = wild type. Values are expressed as means ± SEM. Means without a common letter are statistically different by one-way ANOVA with Tukey–Kramer multiple comparisons post hoc test (P < 0.05).

### Trophoblast-specific inducible *Mtor* knockdown decreases placental TPM SNAT2 and LAT1 expression

To mediate cellular uptake and transplacental transport, the SNAT2 and LAT1 proteins must be translocated to the trophoblast plasma membrane. In addition, membrane trafficking of SNAT2 and LAT1 is tightly regulated in cultured primary human trophoblast cells by mTOR signaling^32^. We used western blotting to determine SNAT2 and LAT1 protein abundance in TPM isolated from WT, *Mtor*, and *Mtor*^kD^ placentas. As shown **in Figure 7 a and b,** TPM SNAT2 (−42%, p=0.006, n=3-4 dams/each group) and LAT1 (−43%, p=0.02, n=3-4 dams/each group) were significantly lower in the *Mtor*^kD^ group as compared to than WT and *Mtor* groups.

**Figure 7:**
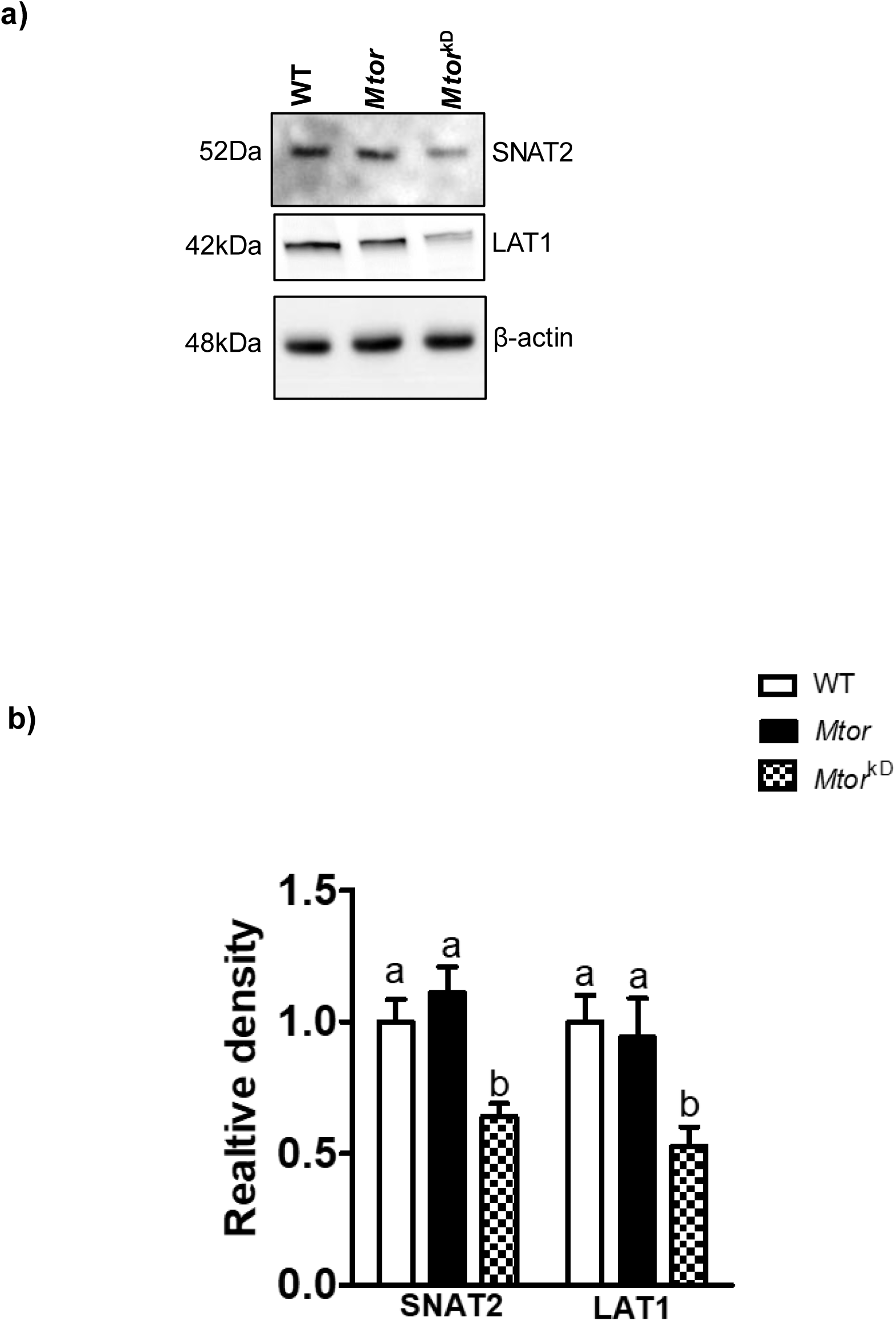
Decreased trophoblast plasma membrane SNAT2 and LAT1 expression following induction of trophoblast-specific mTOR knockdown. mTOR knockdown was induced by the administration of doxycycline starting at E14.5 (*Mtor* ^kD^). In control transgenic animals (*Mtor*), the vehicle was administered at E14.5. At E17.5, animals were sacrificed, and placentas from each litter were pooled and homogenized. Trophoblast plasma membranes were isolated, and the protein expression of the amino acid transporter isoforms SNAT2 (System A) and LAT1 (System L) was determined using Western blot. (a) Representative western blots of SNAT2 and LAT1 expression in TPM of WT, *Mtor,* and *Mtor*^kD^ placentas. Equal loading was performed. (b) Summary of the western blot data. n = 3-4 dams in each group. WT = wild type. Values are expressed as means ± SEM. Means without a common letter are statistically different by one-way ANOVA with Tukey– Kramer multiple comparisons post hoc test (P < 0.05).

### Trophoblast-specific inducible *Mtor* knockdown decreases placental TPM System L and System A amino acid transport

In TPM isolated from *Mtor*^kD^ placentas, the capacity to transport System L (−62%, p=0.002, n=3-4 dams/each group) and System A (−55%, p=0.001, n=3-4 dams/each group) amino acid transport was markedly decreased compared to TPM isolated from WT, or *Mtor* placentas (**Figure 8 a and b**).

**Figure 8:**
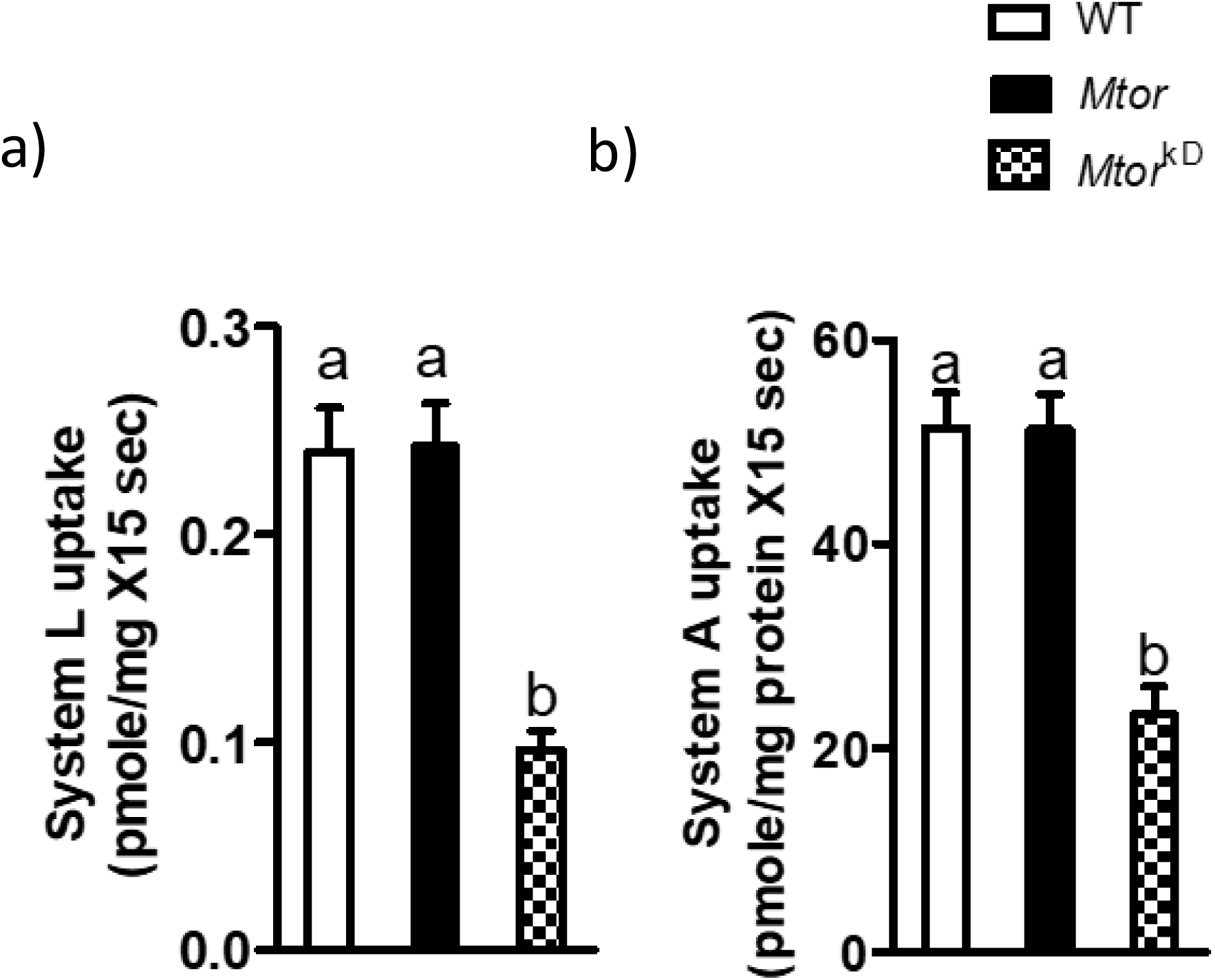
Decreased trophoblast plasma membrane System L and System A activities following induction of trophoblast-specific mTOR knockdown. mTOR knockdown was induced by the administration of doxycycline starting at E14.5 (*Mtor* ^kD^). In control transgenic animals (*Mtor*), the vehicle was administered at E14.5. At E17.5, animals were sacrificed, and placentas from each litter were pooled and homogenized. Trophoblast plasma membranes were isolated. System L (a) and System A (b) transporter activities were determined using isotope-labeled substrates and rapid filtration techniques in TPM isolated from WT, *Mtor,* and *Mtor*^kD^ placenta at E 17.5. Values are expressed as means ± SEM. Means without a common letter are statistically different by one-way ANOVA with Tukey–Kramer multiple comparisons post hoc test (P < 0.05).

## Discussion

Our study represents the first application of piggyBac transposase-enhanced transgenesis to achieve inducible trophoblast-specific gene modulation. In addition, we demonstrate that inhibition of trophoblast mTOR signaling is mechanistically linked to decreased placental nutrient transport and reduced fetal growth.

Despite their introduction to the field many years ago, the successful use of approaches to achieve trophoblast-specific gene modulation has been rather limited. For example, the hCYP promoter-Cre construct is associated with some degree of ‘leakiness’, sometimes resulting in expression of the construct in fetal tissues such as the brain, eye, skin and heart ^64^. Moreover, placental defects are highly prevalent in embryonic lethal mouse mutants ^65^, including *Mtor,* precluding meaningful studies of the function of some critical placental genes in the second half of pregnancy. Herein, we report using piggyBac transposase-enhanced transgenesis to achieve trophoblast-specific gene knockdown. We believe that our approach represents a robust tool for inducible trophoblast-specific gene modulation that will allow the placental research field to conduct detailed mechanistic studies of the function of hundreds of placental genes *in vivo* without risking that global deletion of these genes results in embryonic lethality. In addition, many placental genes regulate both placental development and the function of the mature placenta, requiring experimental models with temporal control of trophoblast-specific gene modulation, such as piggyBac transposase-enhanced transgenesis, to differentiate between these two distinct processes.

Placental mTORC1 and 2 signaling pathways are positive regulators of key amino acid transporters in PHT cells and human placental villous explants ^32,38,66^. Inhibition of mTORC1 and/or mTORC2 downregulates trophoblast System A and L amino acid transporter activity by decreasing the plasma membrane expression of specific System A (SNAT2, *SLC38A2*) and System L (LAT1, *SLC7A5*) isoforms with no effect on overall protein expression ^17^. Distinct mechanisms mediate the regulation of SNAT2 and LAT1 membrane trafficking by mTORC1 and 2. Inhibition of mTORC1 results in activation of Nedd 4-2, an E3 ubiquitin ligase, which ubiquitinates the specific amino acid transporter isoforms leading to their withdrawal from the plasma membrane and degradation^31^. In contrast, inhibition of mTORC2 decreases SNAT2 and LAT1 plasma membrane abundance in PHT cells mediated by downregulation of Rho GTPase Cdc42 and Rac1^30^, which are essential for actin skeleton function. In the current study, trophoblast-specific knock-down of *Mtor*, resulting in a marked inhibition of placental mTORC1 and mTORC2 signaling, decreased the protein expression of SNAT2 and LAT 1 in the trophoblast plasma membrane, providing the first in vivo evidence of mTOR controlling the trophoblast plasma membrane trafficking of these two specific amino acid transporter isoforms. Importantly, using isolated trophoblast plasma membrane vesicles incubated with radiolabeled amino acids and rapid filtration, we confirmed that trophoblast-specific mTOR knockdown causes a marked decrease in System A and L amino acid transporter function. We propose that a decreased transplacental transport of amino acids causes the inhibition of fetal growth following trophoblast-specific mTOR knockdown in our model. This aligns with our recent study demonstrating that placental-specific Slc38a2 knockdown in mice resulted in IUGR ^63^.

Fetal weight was reduced in trophoblast-specific inducible *Mtor* knockout mice, which is similar in magnitude to that observed in other mouse models either with inhibition of placental mTOR signaling due to maternal folate deficiency ^44^, or following administration of rapamycin to pregnant mice ^67^. Our study demonstrates that inhibition of mTOR signaling only in the trophoblast is sufficient to cause IUGR. Using Trophoblast-specific Cre recombinase transgene driven by the CYP19 promoter to knock down placental mTOR, Akhaphong and co-workers previously reported decreased fetal weight in females but not in male pups ^52^. However, the data in this previous study is difficult to interpret given that a conditional gene-targeting approach was not used, making it difficult to differentiate between the effect on placental development and function, neither inhibition of placental mTOR signaling nor specificity of the mTOR knockdown to the trophoblast only was confirmed, and placental function was not assessed ^52^.

Our findings have clear translational significance. First, placental mTOR signaling is inhibited in human IUGR^12,38–41^ and increased in fetal overgrowth associated with maternal obesity and/or GDM^14,49–51,68^. Second, placental System L activity has been reported to be reduced in pregnancies complicated by IUGR^11,69^, associated with a decrease in fetal circulating concentrations of many essential amino acids^70,71^. In addition, SNAT2 (SLC38A2) protein expression and/or System A activity are decreased in term placenta of IUGR or small for gestational age fetuses ^9,10,72,73^. Maternal obesity, which increases the risk of fetal overgrowth, has been reported to be associated with increased placental amino acid transport capacity^14^. Specifically, System A activity, but not System L, was positively correlated with birth weight in a cohort of normal and obese women, and transplacental amino acid transport mediated by System A is increased in a mouse model of maternal obesity and fetal overgrowth^74^. Thus, the mechanistic link between inhibition of placental mTOR signaling, reduced placental amino acid transport activity, and IUGR reported here provides novel insights into the causes of human IUGR.

In the present study, we developed a novel method for achieving inducible trophoblast-specific *Mtor* knockdown *in vivo* in mice, allowing us to generate novel mechanistic information on regulating placental function and fetal growth. We demonstrate that inhibition of trophoblast mTOR signaling in late pregnancy is mechanistically linked to decreased placental nutrient transport and reduced fetal growth. Currently, no specific intervention strategies are available that can be used clinically to treat IUGR and fetal overgrowth. We speculate that placenta-specific targeting of mTOR signaling represents a promising avenue to improve outcomes in pregnancies complicated by abnormal fetal growth. This could be made possible by recent developments in nanotechnology, which have provided an innovative approach for drug delivery to or gene targeting of the placenta^75^.

## Data Availability

All supporting data for this manuscript are included in the Figures.

